# Contextualised real-time mass spectrometry improves glycosylation detection and characterisation

**DOI:** 10.64898/2026.07.03.736344

**Authors:** Maia I. Kelly, Christopher Ashwood

## Abstract

Glycosylation is a structurally diverse, non-template-driven modification whose analysis by liquid chromatography-mass spectrometry is constrained by discovery-mode acquisition rules developed for proteomics. Data-dependent acquisition filters, such as intensity-based precursor selection and charge-state exclusion, map poorly onto glycan analysis, which span wide ranges of charge state and abundance independent of their biological importance. Here we present glycosylation real-time mass spectrometry (GlycoRTMS), an instrument-API method that annotates observed precursor masses with glycan compositions in real time and uses this context to guide fragmentation. Composition-aware precursor prioritisation sampled deeper into the precursor space, expanding MS2 coverage of a hyaluronic acid hydrolysate from four to eight oligosaccharide subunits. Charge-state-specific collision energy equations tailored to oligosaccharides produced complete fragment ladders where fixed normalised collision energy did not. MS3 triggering gated by both diagnostic ions and glycan composition matching enabled efficient, chromatography-compatible characterisation of *O*-acetylated sialic acids and identified product ions specific to *O*-acetylation. Together, these strategies improve both the depth and quality of glycan detection and characterisation within a single injection.

## Introduction

Glycosylation is among the most prevalent and structurally diverse modifications of proteins and lipids. Unconjugated sugars can also act as glycosylation substrates, generating oligosaccharides and polysaccharides. These sugar-based molecules decorate the extracellular matrix and are involved in a range of cellular processes including protein folding, trafficking, immune recognition and pathogen interaction. Unlike the genome or proteome, the glycome is not template-driven, instead it is the result of the combinatorial action of glycosyltransferases and glycosidases to sequentially generate the glycan structures present. This complexity places a heavy burden on glycan analysis, which aims to resolve monosaccharide presence, frequency and how they are connected and modified.

Liquid chromatography–mass spectrometry (LC-MS) serves as an important technique to address this complexity, as mass-based profiling provides insights regarding the monosaccharide composition of the intact molecule. Further, liquid chromatography enables separation of compositions with the same mass for isomeric identification. These molecules can be subjected to fragmentation, which can elucidate the monosaccharide composition of the intact glycan and how the monosaccharides are arranged. At the inception of MS, data would be acquired iteratively, following data analysis and subsequent identification of targets for fragmentation^1,2^, however today, advances made in discovery-mode acquisition methods enable real-time chromatography-compatible data acquisition that reduces the need for repeated injections of the same sample^3–5^.

The dominant methodology for discovery-mode acquisition is data-dependent acquisition (DDA) in which the instrument surveys a full scan (either MS1 or MSn) and selects ions for fragmentation according to user-defined rules, first developed by Yates *et al* in 1995^6^. This approach is now widely applied across the mass spectrometry field, enabling LC-compatible discovery methods to both identify and characterise molecules as they elute from a column. These user-defined rules enhance the specificity and likely successful characterisation rate of the triggered fragmentation scans. The more frequently used rules for bottom-up proteomics include ignoring singly charged ions, dynamic exclusion, and the most widely used, prioritisation of the most intense analytes from each scan. These filters translate poorly to glycan discovery, as glycans can be observed across a wide span of charge states (−1 to beyond −6) and vary by orders of magnitude in abundance, independent of their biological importance.

More recent approaches have shifted the analytical interpretation onto the instrument itself, making fragmentation decisions in real time using contextual knowledge unavailable to standard vendor logic. The development and accessibility of instrument application programming interfaces (IAPIs) enabled this possibility, and advancement has been largely localised to the proteomics community, producing a succession of methods that go beyond existing vendor method offerings. The Gygi and Schweppe laboratories, in particular, advanced real-time adaptive control through TOMAHAQ^7^ and Real Time Library Search^8^, respectively. Through the assignment of peptide context to observed *m/z* values, both approaches improve the identification and quantitation of peptides.

To date, these advanced real-time approaches have been applied almost exclusively to the field of proteomics, leaving glycans and glycopeptides, one of the more challenging analyte classes, dependent on non-optimal vendor filters. Here we adapt the principle of real-time, identity-aware acquisition to glycomics. We describe glycosylation real-time mass spectrometry (GlycoRTMS) which annotates observed *m/z* values with glycan compositions, providing a glycosylation context to observed analytes prior to fragmentation. GlycoRTMS enables strategies that are inaccessible to context-independent vendor methods. The presented method utilises prioritisation of precursors that match glycan compositions rather than relying on intensity, and charge-state-specific collision energy equations tailored to oligosaccharides. Additionally, when a precursor is matched to a glycan composition and a structurally diagnostic product ion is observed in the MS2 spectrum, MS3 can be triggered. We show that these capabilities improve both the depth and quality of glycan detection and characterisation within a single LC-MS injection.

## Results and discussion

At the initialisation of every LC-MS run using GlycoRTMS, GlyCombo generates a glycan composition list from user-specified parameters based on the sample under study. An LC-MS run focused on mammalian *N*-glycan context, for example, matches only neutral masses consistent with the monosaccharides present in mammalian *N*-glycosylation **(Figure 1A)**. During the run, each MS1 spectrum is deisotoped and transformed into a list of neutral monoisotopic masses, which are matched against the GlyCombo composition list to provide glycosylation context for the observed precursors. Dynamic exclusion is built into the program to minimise redundant MS2/MS3 scans. Matched precursors are then prioritised by intensity, and the instrument is triggered to perform MS2 on the corresponding precursor *m/z* values. An optional MS3 event can subsequently be triggered based on the presence of diagnostic product ions. By supplying this glycosylation context, GlycoRTMS enables several real-time strategies: precursor prioritisation, glycan-specific fragmentation parameters, and MS3-based enhanced characterisation **(Figure 1B)**.

**Figure 1.**
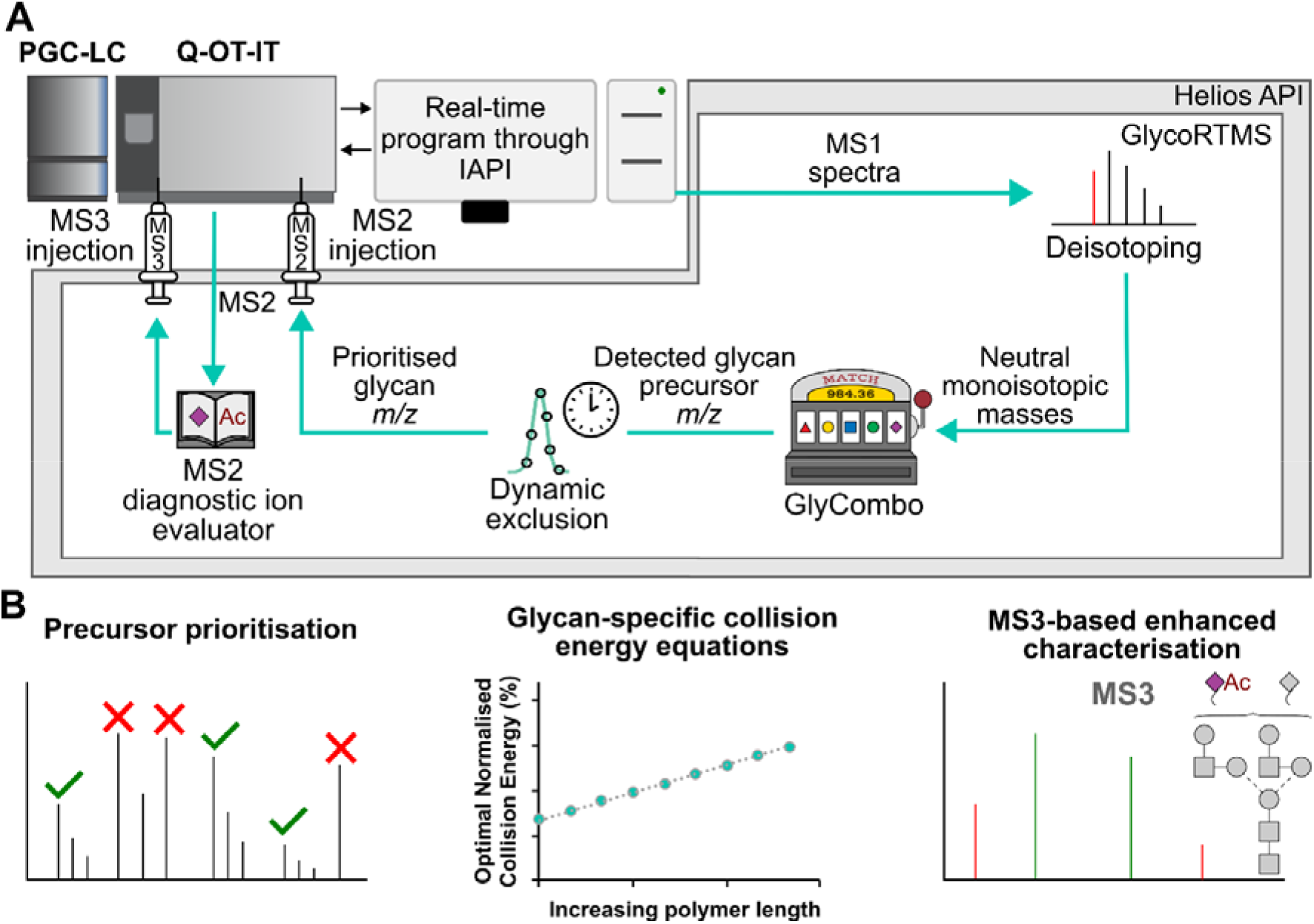
GlycoRTMS contextualises real-time decision-making with glycan-specific context. **A**The real-time assignment of glycan compositions from GlyCombo to MS1 precursors enables real-time MS2 and MS3 scans. **B** Outcomes achieved by real-time contextualisation of observed glycans, including precursor prioritisation, glycan-specific collision energy equations, and informed MS3 triggers

To evaluate GlycoRTMS glycan prioritisation, partially acid-hydrolysed hyaluronic acid was analysed by a standard method and a GlycoRTMS-enabled method. These methods were identical with the exception of the glycan composition matching component employed with GlycoRTMS **(Figure 2A)**. When the glycan matches were plotted in the context of a standard LC-MS run, glycan compositions spanned both the highest- and lowest-intensity MS2 scans, with approximately two orders of magnitude separating the most and least intense, while non-glycan ions (*e.g*. chemical and background noise) accounted for approximately half of all MS2 scans **(Figure 2B)**. Glycan prioritisation increased the maximum MS2 intensity and expanded MS2 coverage of the hyaluronic acid hydrolysate from four polymers (2, 3, 4, and 6 repeats) to eight polymers (2–10, excluding 9) **(Figure 2C)**. Despite their low intensity, which would have caused them to be unsampled by standard DDA, these subunits yielded full linear sequence coverage in the MS2 scan **(Figure 2D)**. Together, these results demonstrate that GlycoRTMS-enabled prioritisation triggers MS2 on lower intensity glycans from MS1 scans, achieving sensitive glycan detection against noisy backgrounds. Having established improved MS2 triggering, we next sought to use the glycan context to improve MS2 quality.

**Figure 2.**
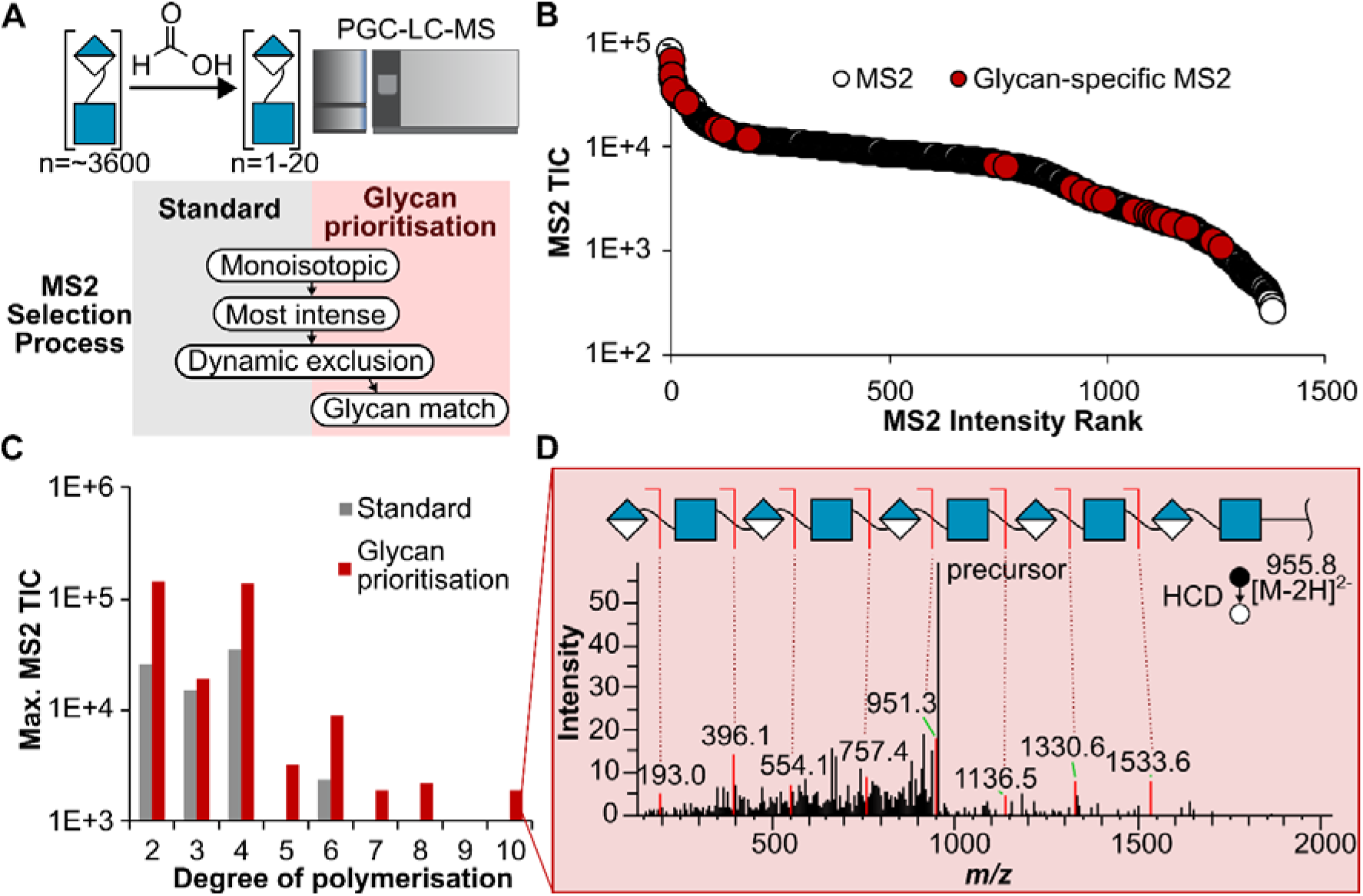
Real-time glycan assignment improves MS2 depth through prioritisation of precursors assigned to hyaluronic acid subunits. **A** Glycan prioritisation approach for acid hydrolysed hyaluronic acid. **B** Intensity rank distribution of glycans compared to background, across an LC-MS run. **C-D** Glycan prioritisation enables detection and characterisation of less intense, larger hyaluronic acid subunits

A principal goal of MS2 fragmentation is to dissociate the analyte and decipher the connectivity of its constituent monosaccharides and chemical groups^9^. To make a single collision energy setting applicable across diverse molecules, normalised collision energy (NCE) approaches have been developed. On ThermoFisher Scientific instruments, this value is calculated by determining the minimum calibrated collision energy required to fully dissociate three calibrants (caffeine, MRFA peptide, and Ultramark 1621) spanning a range of *m/z*; plotting the optimal collision energy of each calibrant against its *m/z* yields an instrument-specific equation expressed as NCE^10^. Because this calibration does not use carbohydrates or multiply-charged analytes, it presents a clear opportunity for improvement through a glycosylation-specific context. Colominic acid, a sialic acid (Neu5Ac) polymer found in the bacterial capsule, carries a carboxylic acid that often retains a negative charge, making it an ideal proof of concept for *m/z*- and charge-specific fragmentation optimisation. A targeted LC-MS acquisition was performed on Neu5Ac repeats of 2–9 across deprotonated ([M-nH]^-n^) charge states −1 to −4, and the data were imported into Skyline for interpretation **(Figure 3A)**. For each species, the optimal collision energy was defined as that producing the greatest MS2 total ion current (TIC), and these values were used to construct lines of best fit **(Figure 3B)**. Each charge state exhibited distinct slopes and intercepts, underscoring the need for charge-state-specific corrections; the exception was singly charged glycans, which performed best at a fixed NCE. The resulting slopes and intercepts were applied to the GlyCombo output to inform a GlycoRTMS acquisition.

**Figure 3.**
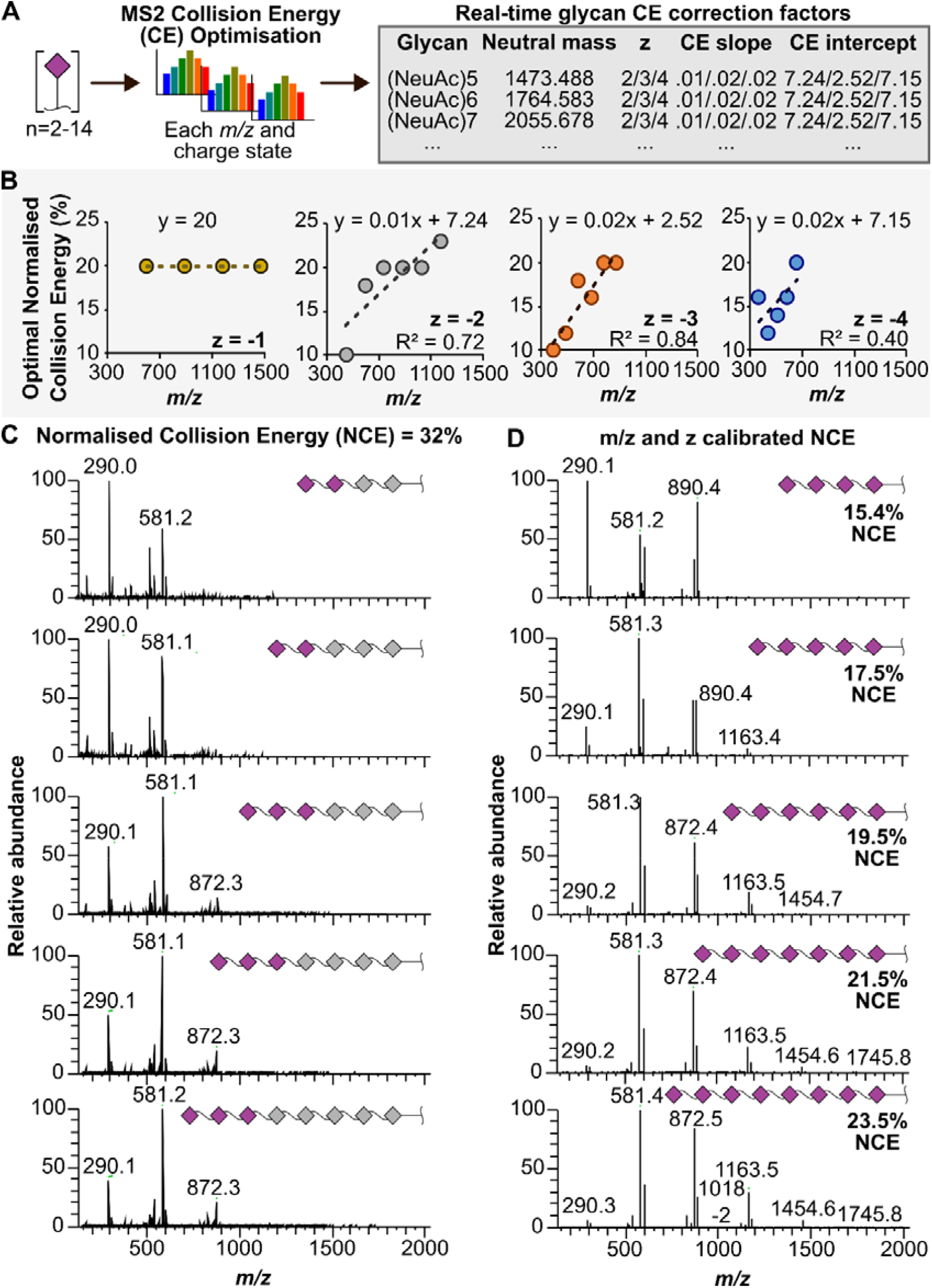
Collision energy adjustment improves glycan coverage compared to fixed NCE. **A** Optimisation of Neu5Ac polymer optimal collision energy to develop CE correction factors for real-time MS. **B** Charge state-specific collision energy optimisation curves reveal standard normalised collision energy is unsuitable for multiply-charged oligosaccharides. **C** Fixed NCE results in insufficient oligosaccharide fragment coverage. **D** Corrected NCE results in complete fragment series for oligosaccharides.

A control GlycoRTMS acquisition, with glycan prioritisation but without collision energy (CE) correction, used a fixed NCE (Figure 3C). MS2 coverage was incomplete for all sialic acid polymers, with a maximum of three sialic acids covered by the fragment series at 32% NCE; moreover, the MS2 spectra for increasing polymer lengths were near-identical, so glycan composition could only be assigned by reference to precursor *m/z* and charge. By contrast, the variable-NCE approach encoded in the GlyCombo output achieved full sequence coverage while minimising intact precursor **(Figure 3D)**. Compositions beyond those used to derive the CE correction equations also produced high-quality MS2 spectra **(Supplementary Figure 2)** demonstrating the extensibility of the approach. These results demonstrate the importance of appropriately tailored collision energy regimes, as has likewise been shown in bottom-up proteomics^11^.

Thus far we have shown that GlycoRTMS achieves deeper MS2 sampling of glycans, enabling characterisation of species that would have been unobserved by non-real-time methods. MS3-based methods are inherently slower, since the additional isolation and fragmentation events substantially reduce ion current, but they yield further structural information about the specific feature isolated for MS3^12^. To offset this speed penalty, we used intelligent prioritisation to trigger MS3 in real time only on the appearance of the *O*-acetylated Neu5Ac diagnostic ion for precursors matching a glycan composition list (Figure 4A, *m/z* 332). As a result, MS3 spectra contained only fragments derived from *O*-acetylated sialic acid, improving confidence in the structural specificity of the MS2 productions.

**Figure 4.**
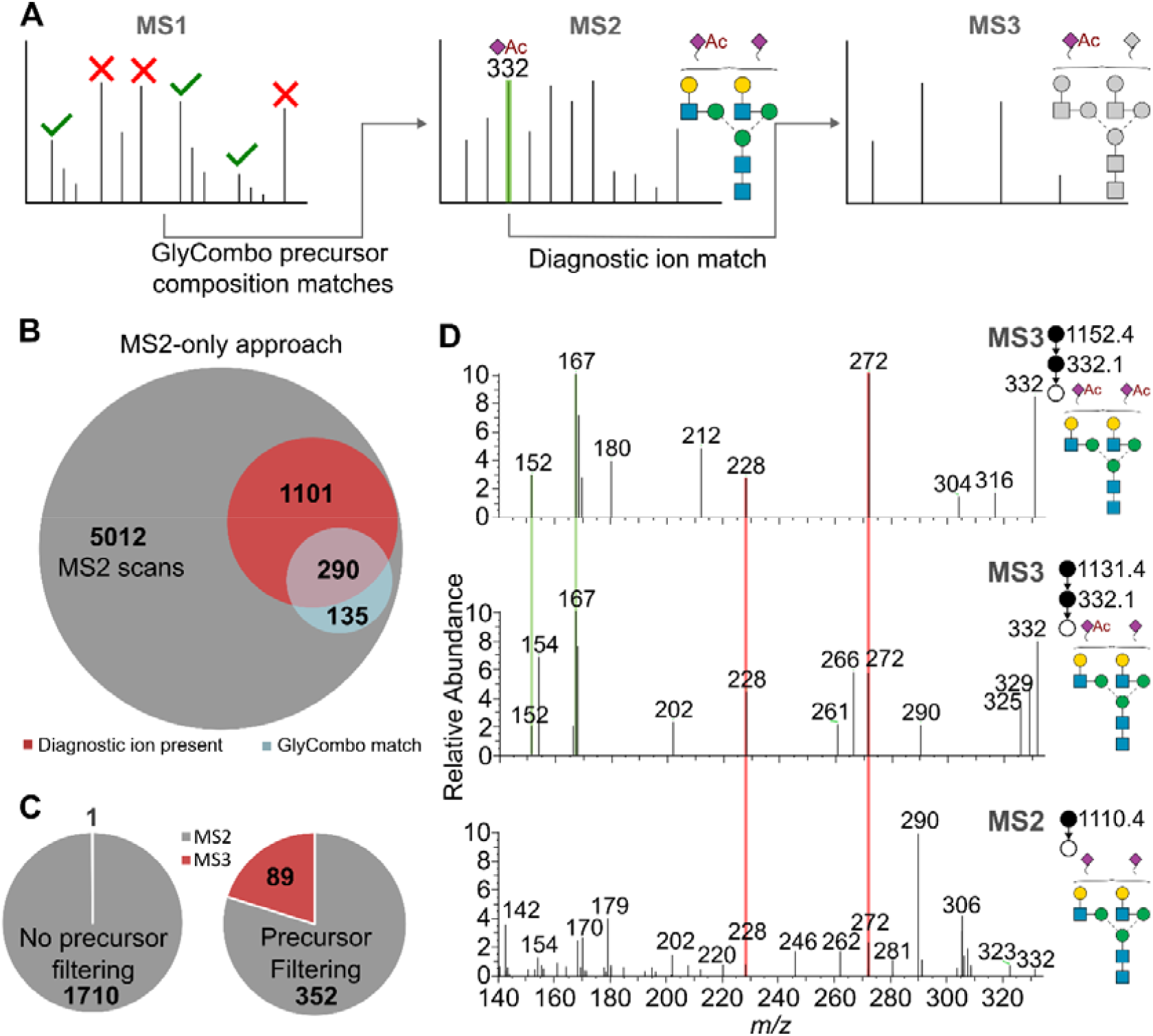
Real-time triggering of MS3 enables efficient, chromatography-compatible glycan structural analysis. **A** Overall workflow for real-time MS3 triggering with an *O*-acetylated Neu5Ac diagnostic ion. **B** A small subset of MS2 scans satisfy both correct precursor mass and diagnostic ion presence. **C** Precursor filtering to glycans is required to ensure MS3 triggering on *O*-acetylated glycans. **D** Real-time MS3 identifies unique diagnostic ions for *O*-acetylation.

Building on our previous work on rat serum *N*-glycosylation^13^, standard methods require both a glycan precursor match and the presence of a compositionally informative ion to constitute evidence of *O*-acetylation **(Figure 4B)**. Following the same logic, the GlycoRTMS MS3 triggers required both a glycan composition match to the precursor and detection of the diagnostic ion. Without precursor filtering for *N*-glycan compositions, only a single MS2 scan containing the *m/z* 332 compositional ion was sampled, resulting in one MS3 event. Precursor filtering reduced the total number of MS2 scans by 80% yet produced 89 MS3 triggers, demonstrating that filtering was necessary to select glycan precursors with the potential to be O-acetylated (Figure 4C). With this refined approach, we could evaluate diagnostic ion specificity in real time.

Comparing MS3 spectra across structures with no, partial, and complete O-acetylation revealed two product ions (*m/z* 152 and 167) present only in the *O*-acetylated structures **(Figure 4D)**. In contrast, *m/z* 332, 272, and 228 were all detected in the un-*O*-acetylated *N*-glycan MS2 and were therefore designated non-specific. The real-time MS3 method thus identified *m/z* 152 and 167 as specific product ions produced by *O*-acetylation of Neu5Ac. As a proof of concept, this strategy can be extended to evaluate diagnostic ions for any glycan structural feature, such as phosphorylation or sulfation, provided that glycans with varying occupancy are detected, and could in principle resolve specific linkages (e.g. 7-, 8-, or 9-O-acetylation).

## Conclusions

By supplying real-time glycan-composition context to the mass spectrometer, GlycoRTMS overcomes key limitations of standard discovery-mode acquisitions for glycans. Prioritising precursors that match glycan compositions rather than raw intensity, applying oligosaccharide-tailored collision energies, and gating MS3 on structurally diagnostic product ions together improve the depth, quality, and structural informativeness of glycan analysis within a single LC-MS injection. The approach presented here is identity-aware rather than tied to any one glycan class, and is able to generalise beyond the systems demonstrated here. The same diagnostic-ion logic could resolve other labile or isomeric features, such as phosphorylation, sulfation, or specific *O*-acetylation linkages, provided appropriate composition lists and diagnostic ions are supplied. More broadly, GlycoRTMS extends the real-time, identity-aware acquisition strategies established in proteomics to the glycosciences.

## Methods

### Chemicals, glycan sources and preparation

All chemicals and reagents were purchased from Millipore Sigma (Castle Hill, Australia) unless specified otherwise. Hyaluronic acid (53747) was hydrolysed at 80 °C for 20 min in pure formic acid.

*N*-glycans were released from rat serum as previously described^13^, briefly, rat serum was solubilised in 5% sodium dodecyl sulfate/100 mM triethylammonium bicarbonate. Proteins were reduced (5 mM dithiothreitol, 55 °C, 30 min), and alkylated (10 mM iodoacetamide, RT, 30 min, dark). Glycans were released and purified for LC-MS as in described in Ashwood *et al*.^14^: proteins were precipitated, immobilised on DNA miniprep silica columns (Bioneer), and N-glycans liberated with 5 units PNGase-F (Promega) in 50 mM TEAB (pH 8) at 37 °C overnight, followed by elution with 400 μL 0.1% formic acid.

Colominic acid (C5762), released serum *N*-glycans, and the acid hydrolysate of hyaluronic acid were brought up to 400 μL with an aqueous 0.1% formic acid solution and desalted on Agilent Carbon-S SPE (100 mg): conditioned with acetonitrile/0.1% formic acid, equilibrated with water/0.1% formic acid, loaded with sample, washed with water/0.1% formic acid, and eluted with 50:50 acetonitrile:water/0.1% formic acid. Eluates were dried by centrifugal evaporation, resuspended in ultra-pure water, and transferred to a 96-well PCR plate for injection.

### LC-MS setup

Glycans were separated with a Thermo Fisher Scientific Vanquish Horizon HPLC (San Jose, USA) and ionised into an Orbitrap IQ-X Tribrid mass spectrometer (San Jose, USA). A Merck Millipore Supel Carbon LC column (150 mm length by 1 mm internal diameter, 2.7-micron pore size), held at 80 °C, was used for all separations. To achieve supercharging^15^ and collapse non-reduced glycan anomers^16^, mobile phase A was composed of water and mobile phase B was composed of acetone with 5 mM hexafluoroisopropanol and 5 mM butylamine added. Glycans were separated over a 20 min run, with 0-30% B over 9 min, 100% B held for 4 min, then 100% A for 4 min.

A Thermo Scientific IQ-X Tribrid was set to negative mode with a total cycle time of 3 seconds. ESI voltage was 3.5 kV, with sheath and auxiliary gases at 30 and 10 arbitrary units, respectively. Precursor spectra were collected across a full scan of 340–1700 *m/z* in an Orbitrap at 240,000 resolution to hit an AGC target of 1e^6^ with a maximum injection time of 502 ms.

### Standard comparison Methods

From the acquired MS1 scan, DDA MS2 was performed with dynamic exclusion filtering: exclusion duration of 8 seconds, 0.75 *m/z* mass tolerance low and high, and excluding isotopes. All MS2 were performed with beam-type CID (referred to as HCD) with the following parameters: isolation window of 1.6 m/z, 32% NCE, 200 ms maximum injection time, and AGC target of 1e^4^.

### Real-time methods

The IAPI program, GlycoRTMS, was developed in C# using the Helios^17^ package (v1.0.0.13, compiled from source). For each MS1 scan acquisition, peaks were extracted as *m/z*, intensity, and charge values, then deisotoped using a charge-aware algorithm to only retain monoisotopic peaks. Deisotoped, charge-assigned precursors were converted to neutral masses and matched against a glycan composition database generated by GlyComboCLI^18,19^ at the start of each injection. Optionally, these GlyComboCLI export files contained charge-state-dependent collision energy calibration parameters, and diagnostic ion *m/z* values. All injection and confirmation events were written to an execution log for downstream performance evaluation.

MS2 and MS3 scans were injected into the mass spectrometer after IAPI-based dynamic exclusion filtering: exclusion duration of 6 seconds, 0.75 *m/z* mass tolerance low and high, and excluding isotopes. All MS2 were performed with HCD with the following parameters: isolation window of 2 *m/z*, variable NCE, 200 ms maximum injection time, and AGC target of 1e^4^. All MS3 were performed with HCD with the following parameters: isolation window of 2 *m/z*, 40% NCE, 200 ms maximum injection time and AGC target of 1e^4^.

### Collision energy optimisation

A shortened LC-MS method was used for collision energy optimisation with glycans separated over a 14 min run, with 0-30% B over 6 min, 100% B held for 4 min, then 100% A for 4 min. MS1 parameters were identical to the earlier described LC-MS setup. All MS2 were performed with HCD with the following parameters: isolation window of 1.6 *m/z*, variable NCE, 25 ms maximum injection time, and AGC target of 1e^4^. A targeted method was developed using Skyline-daily^20,21^ (version 26.1.1.159) to generate two targeted methods for MS2 collection, with the first injection at 20, 23, 26, 29, and 32 NCE, and the second injection at 10, 12, 14, 16, and 18 NCE. The optimal CE for each *m/z* and z pair was determined by the greatest total ion current for the full scan MS2.

### Data analysis

Glycan composition assignment was performed with GlyComboCLI^18^ and fragmentation spectra were verified and annotated with GlycoWorkBench 2.1^22^.

## Acknowledgements

This research was facilitated by access to Sydney Mass Spectrometry, a core research facility at the University of Sydney. The authors thank Sebastien Gallien and Jean-Jacques Dunyach from Thermo Finnigan for access to the IAPI.

## Data and code availability

Raw LC-MS files are available on GlycoPOST (accession currently unavailable). Skyline CE optimisation chromatograms and GlyCombo output are available on Panorama (https://panoramaweb.org/glycoRTMS.url). In compliance with the usage agreement for the IAPI, pseudocode describing the GlycoRTMS program is provided in Supplementary Figure 1. GlyComboCLI is open-source and available at https://zenodo.org/records/20063129.

## Conflicts of interest

C.A. is the director of Protea Glycosciences, a company which provides fee-for-service glycomics assays, analytical standards, and software.

## Supplementary Figures

**Supplementary Figure 1.**
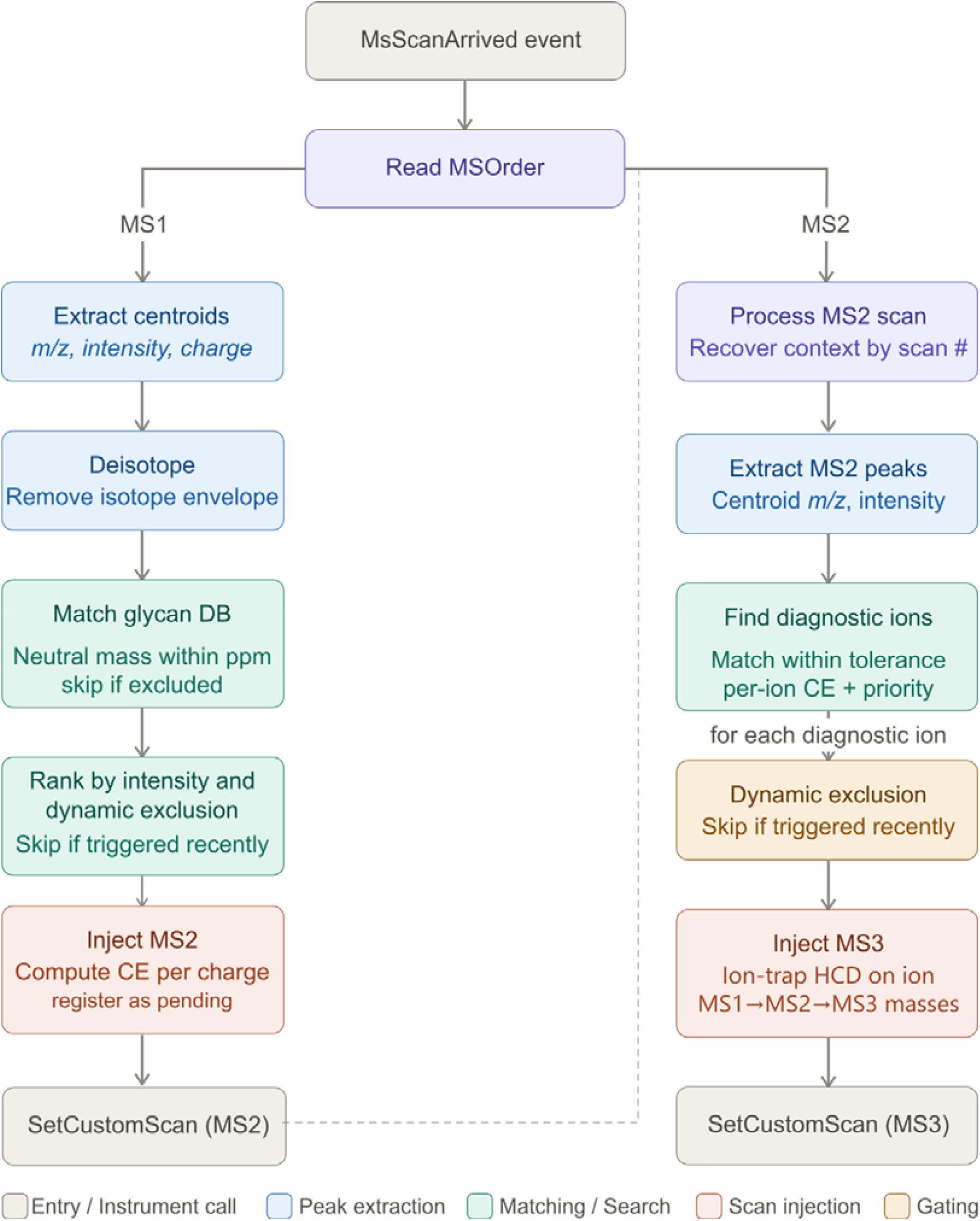
Flow chart describing GlycoRTMS approach

**Supplementary Figure 2.**
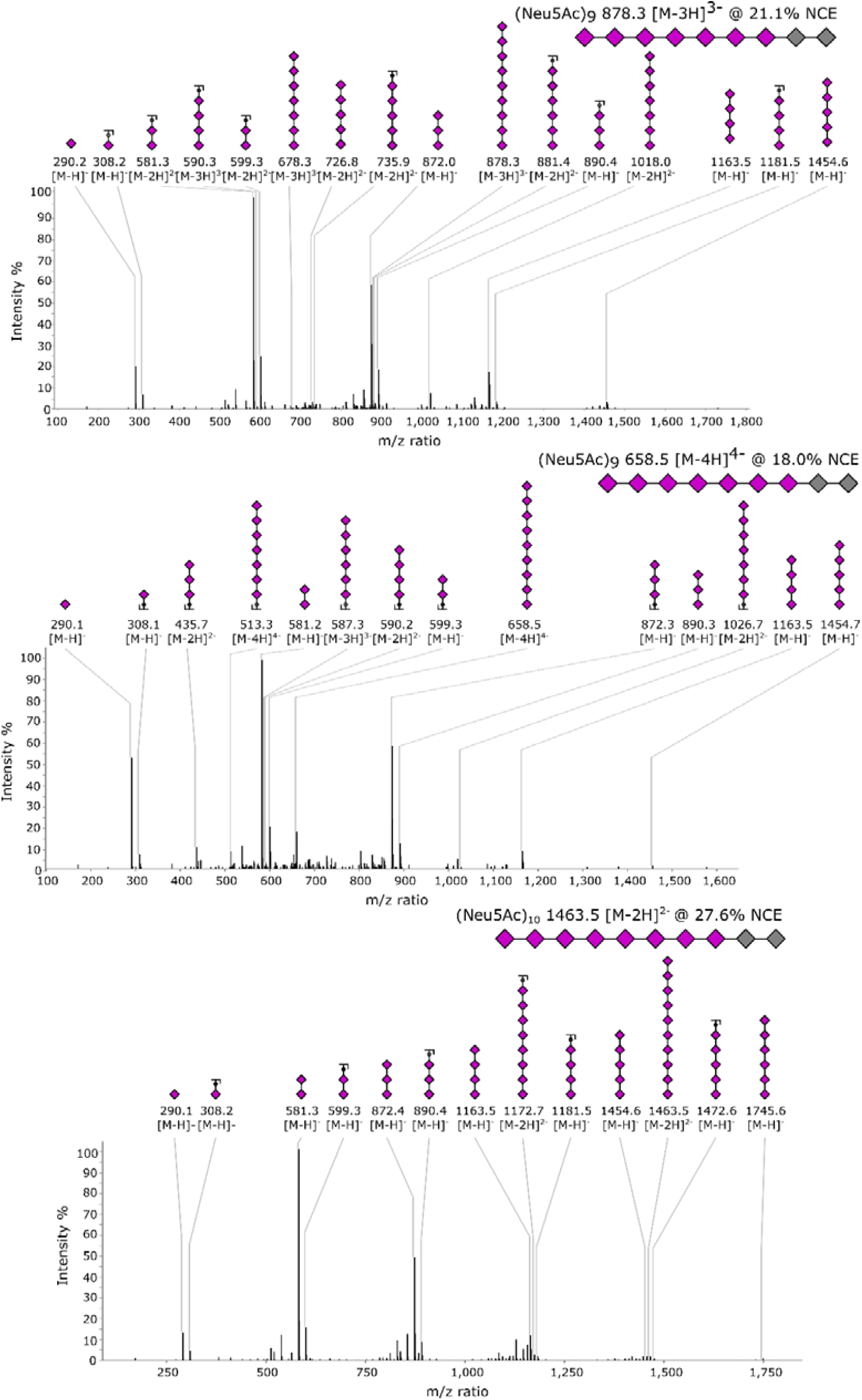
Evaluation of MS2 coverage for colominic acid oligosaccharides beyond the CE optimisation routine

## Notes

https://panoramaweb.org/glycoRTMS.url

