## Supplementary for "Contextualised real-time mass spectrometry improves glycosylation detection and characterisation"

### Supplementary Figures

**Supplementary Figure 1** Flow chart describing GlycoRTMS approach

**Supplementary Figure 2** Evaluation of MS2 coverage for colominic acid oligosaccharides beyond the CE optimisation routine

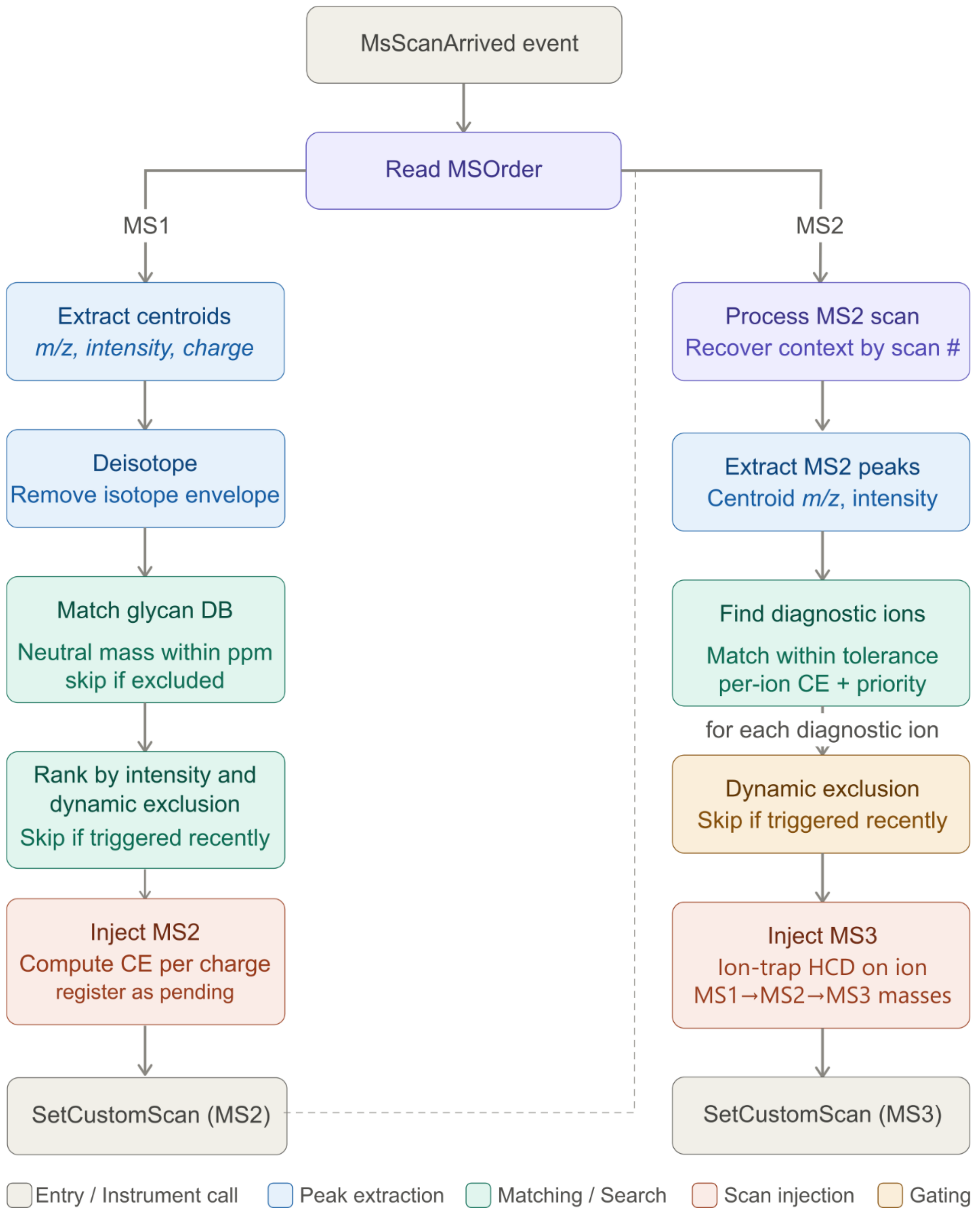


**Supplementary Figure 1** Flow chart describing GlycoRTMS approach

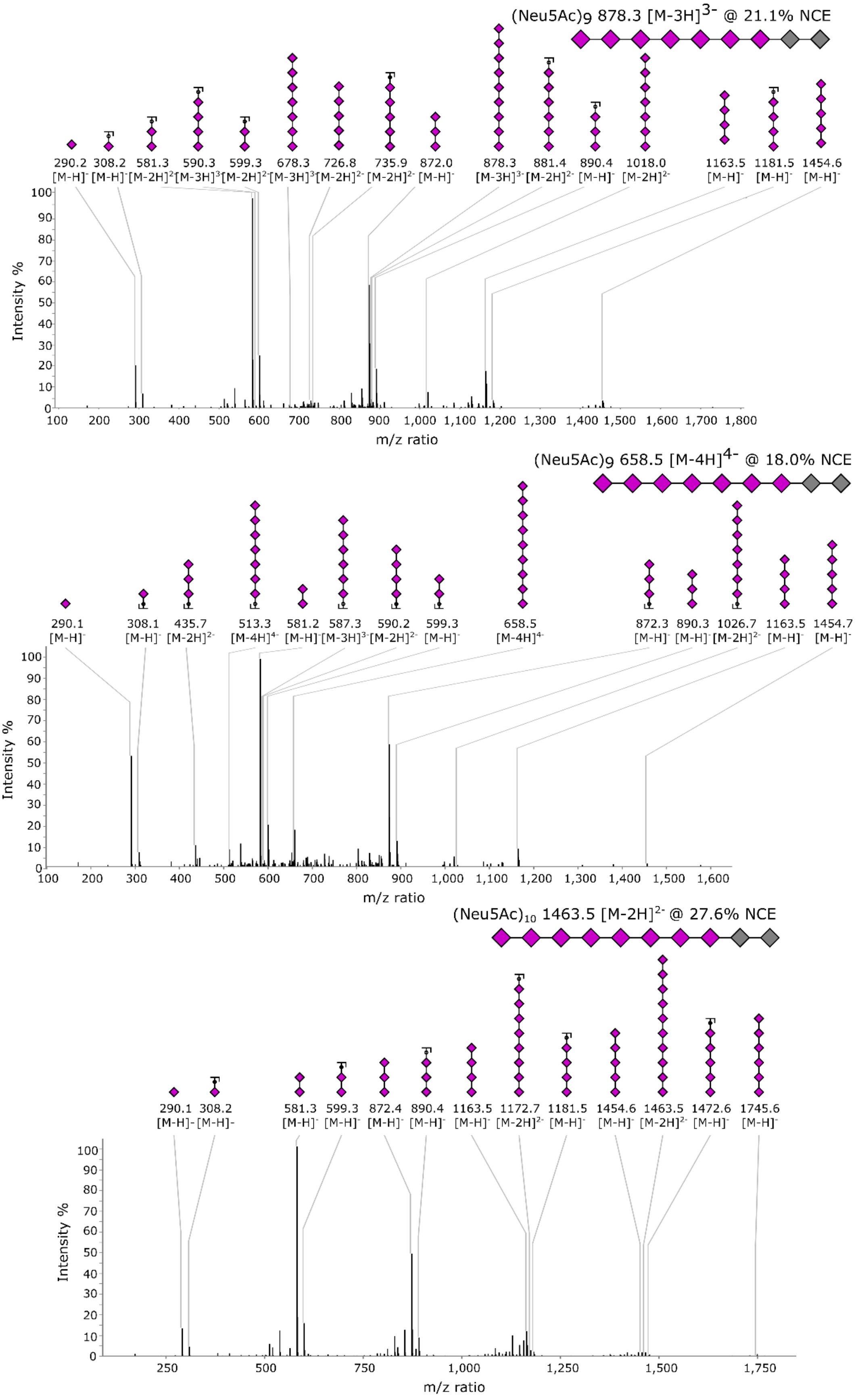


**Supplementary Figure 2** Evaluation of MS2 coverage for colominic acid oligosaccharides beyond the CE optimisation routine
